# Genomic Bayesian confirmatory factor analysis and Bayesian network to characterize a wide spectrum of rice phenotypes

**DOI:** 10.1101/435792

**Authors:** Haipeng Yu, Malachy T. Campbell, Qi Zhang, Harkamal Walia, Gota Morota

## Abstract

With the advent of high-throughput phenotyping platforms, plant breeders have a means to assess many traits for large breeding populations. However, understanding the genetic interdependencies among high-dimensional traits in a statistically robust manner remains a major challenge. Since multiple phenotypes likely share mutual relationships, elucidating the interdependencies among economically important traits can better inform breeding decisions and accelerate the genetic improvement of plants. The objective of this study was to leverage confirmatory factor analysis and graphical modeling to elucidate the genetic interdependencies among a diverse agronomic traits in rice. We used a Bayesian network to depict conditional dependencies among phenotypes, which can not be obtained by standard multitrait analysis. We utilized Bayesian confirmatory factor analysis which hypothesized that 48 observed phenotypes resulted from six latent variables including grain morphology, morphology, flowering time, physiology, yield, and morphological salt response. This was followed by studying the genetics of each latent variable, which is also known as factor, using single nucleotide polymorphisms. Bayesian network structures involving the genomic component of six latent variables were established by fitting four algorithms (i.e., Hill Climbing, Tabu, Max-Min Hill Climbing, and General 2-Phase Restricted Maximization algorithms). Physiological components influenced the flowering time and grain morphology, and morphology and grain morphology influenced yield. In summary, we show the Bayesian network coupled with factor analysis can provide an effective approach to understand the interdependence patterns among phenotypes and to predict the potential influence of external interventions or selection related to target traits in the interrelated complex traits systems.

## Introduction

A primary objective in plant breeding is the develop high yielding varieties with specific grain qualities, resilience to pests and abiotic stresses, and superior adaption to the target environment. As a result, plant breeders devote considerable resources to extensive phenotypic evaluation of germplasm and select on multiple traits. These traits are often correlated at a genetic level through common genetic effects (e.g. pleiotropy) or linkage disequilibrium between quantitative trait locus (QTL). Since multiple phenotypes may exhibit mutual relationships, knowledge of the interdependence among agronomically important traits can improve the efficacy of selection and rate of genetic improvement in systems with complex traits.

In a standard quantitative genetic analysis, multivariate phenotypes can be modeled through multi-trait models (MTM) of Henderson and Quaas (1976) or some genomic counterparts (e.g., Calus and Veerkamp, 2011; Jia and Jannink, 2012) by leveraging genetic or environmental correlations among traits. In particular, MTM has been useful in deriving genetic correlations and enhancing the prediction accuracy of breeding values for traits with low heritability or scarce records via joint modeling with one or more genetically correlated, highly heritable traits (Mrode, 2014). Conventional MTM strategies may provide important insight into the genetic relations between agronomically important traits, but they fail to explain how these traits are related. For instance, consider a case where we have three genetically correlated traits: *y*_1_, *y*_2_, and *y*_3_. With MTM, we cannot address whether the relationship between *y*_1_ and *y*_3_ is due to direct effects, or if the relationship is driven by indirect effects mediated by *y*_2_. Bayesian Networks (BN) offer an effective approach to elucidate the underlying network structure in multivariate data and infer network relationships between correlated variables. A BN is a probabilistic graphical model that represents conditional dependencies among a set of variables via a directed acyclic graph (DAG) (Neapolitan et al., 2004). In the DAG, the variables are represented by nodes, while their conditional dependencies between nodes are indicated with directed edges. In the context of plant breeding, BN can used to elucidate the interdependencies among traits and inform selection decisions for simultaneously improving multiple traits. For instance in the latter case above (*y*_1_ → *y*_2_ → *y*_3_), selection directly on *y*_2_ will affect the quantity of *y*_3_ without an effect on *y*_1_.

With the advent high throughput phenotyping platforms, plant breeders have been provided with a suite of tools for phenotypic evaluation of large populations (Shakoor et al., 2017). These platforms leverage robotics, precise environmental control, and remote sensing techniques to provide accurate, repeatable and high resolution phenotypes for large breeding populations throughout the growing season (Araus and Cairns, 2014; Shakoor et al., 2017; Araus et al., 2018). These data can be used to redefine characteristics underlying superior agronomic performance by quantifying secondary traits associated with seedling vigor, plant architecture, photosynthesis, transpiration, disease resistance, and stress tolerance (Cabrera-Bosquet et al., 2016; Sun et al., 2017; Crain et al., 2018). However given these new approaches, breeders are faced with the new challenge of efficiently utilizing these large multidimesional data sets to improve selection efficiency. The primary challenges associated with multivariate analysis and BN approaches using HTP data is that robust parameter estimates can be untenable because the number of estimated parameters within the model increases with the increasing number of phenotypes. Moreover even in cases where MTM or BN can be applied, interpreting of interrelationships among a large number of phenotypes can be difficult.

One approach to characterize high-dimensional phenotypes is by using factor analysis (FA). The central idea of FA approaches is to reduce the dimensions of multivariate data sets by constructing unobserved, latent factors, or modules, from correlated phenotypes (de los Campos and Gianola, 2007). The biological importance of these latent factors can be interpreted by inspecting the phenotypes that contribute to each factor. Thus, the advantage of FA for large, multivariate data sets is two fold. First, FA provides a means to reduce the dimensions of multivariate data sets thereby providing statistically sound parameter estimates, and easing visualization and interpretation. Secondly, the latent variables/factors themselves may be representative of underlying biological processes that cannot be observed or measured in the population. For instance, several studies have highlighted the effects of plant hormones such as GA on multiple morphological attributes (Wang and Li, 2006; Lo et al., 2008; Umehara et al., 2008; Bhattacharya et al., 2010; Brewer et al., 2013; Zhou et al., 2013). Thus, a latent factor constructed from these morphological traits may provide information on the biosynthesis or sensitivity of these hormones for individuals within the population. If a certain amount of knowledge regarding the biological role of the variables is already known, a varaint of FA, confirmatory factor analysis (CFA), can be used to estimate latent variables based on predetermined biological classes of observed traits (Joreskog, 1969). These latent variables underlie observed phenotypes and can be evaluated for how well the data support the hypothesis. For instance, Peñagaricano et al. (2015) performed CFA in swine to derive five latent variables from 19 phenotypic traits and inferred BN structures among those latent variables, thereby demonstrating the potential of this approach.

This study aimed to leverage CFA and graphical modeling to elucidate the genetic interdependencies among traits typically recorded in breeding programs (e.g., yield, plant morphology, phenology, and stress resilience). First, we constructed latent variables, using prior biological knowledge obtained from the literature. Then we connected the observed highdimensional phenotypes with these to establish latent variables via Bayesian confirmatory factor analysis (BCFA) to reduce the dimensions of the dataset. Further, factor scores computed from BCFA were considered new phenotypes for a Bayesian multivariate analysis to separate breeding values from noise. This was followed by adjustment of breeding values via Cholesky decomposition to eliminate the dependencies introduced by genomic relationships. Finally, the adjusted breeding values were considered inputs to assess the causal network structure between latent variables by conducting a Gaussian BN analysis. This study is the first, to our knowledge, in rice to characterize various phenotypes with graphical modeling such as BCFA and BN.

## Materials and Methods

### Phenotypic and genotypic data

The rice dataset comprised *n* = 374 accessions sampled from six subpopulations: temperate japonica (92), tropical japonica (85), indica (77), aus (52), aromatic (12), and admixture of japonica and indica (56) (Zhao et al., 2011). The improvement status of each accession was obtained from the USDA-ARS Germplasm Resources Information Network. We used *t* = 48 phenotypes and data regarding 44,000 single-nucleotide polymorphisms (SNP). After removing SNP markers with minor allele frequency less than 0.05, 374 accessions and 33,584 markers were used for further analysis. Of those, 27 phenotypes were reported in Zhao et al. (2011) and McCouch et al. (2016). These phenotypes can be classified into four categories: flowering time (flowering time at three locations, photoperiod sensitivity), grain morphology (seed length, seed width, seed surface area, seed length to width ratio, seed volume), plant morphology (culm habit/angle, flag leaf length and width, plant height at maturity), and yield traits (panicle fertility, seed number per panicle, number of primary branches on the main panicle, panicle length, and the number of panicles on each plant). Zhao et al. (2011) evaluated flowering time-related traits using data from three locations, while the remaining traits were evaluated at one location (Arkansas). The remaining phenotypes were assessed from the salinity stress experiments conducted in Campbell et al. (2017a). These traits were classified into three categories: morphological salt response, ionic components of salt stress, and plant morphology. The class morphological salt response represents how plant growth is affected by salinity stress and is composed of the ratio of shoot biomass of salt stressed plants to control, the ratio of root biomass of salt stressed plants to control, the ratio of the number of tillers for salt stressed plants to control, and two metrics that represent the ratio of shoot height of salt stressed plants to control. Ionic components of salt stress is composed of traits that quantify ions important for salinity tolerance (Na^+^ and K^+^) in both root and shoot tissues. Morphology traits are those that describe the growth of the plant in both control and saline conditions (e.g. shoot biomass, root biomass, shoot height, and tiller number). The data used from Campbell et al. (2017a) were derived from three to six independent greenhouse experiments performed between July and October 2013. Information for all experiments were combined and best linear unbiased estimators were calculated for each line as described in Campbell et al. (2017a). The detailed descriptions of the phenotypes are summarized in Supplementary Table S1.

### Bayesian confirmatory factor analysis

A CFA under the Bayesian framework was performed to model 48 phenotypes. The number of factors and the pattern of phenotype-factor relationships need to be specified in BCFA prior to model fitting. We constructed six latent variables (*q* = 6) from previous reports (Acquaah, 2009; Zhao et al., 2011; Campbell et al., 2017a). The six latent variables derived from our analysis represent the grain morphology, morphology, flowering time, ionic components of salt stress, yield, and morphological salt response (Table S1). Each latent variable captures common signals spanning genetic and environmental effects across all its phenotypes. The latent variables, which determine the observed phenotypes can be modeled as

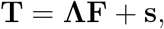

where **T** is the *t × n* matrix of observed phenotypes, **Λ** is the *t × q* factor loading matrix, **F** is the *q × n* latent variables matrix, and **s** is the *t × n* matrix of specific effects. Here, **Λ** maps latent variables to the observed variables and can be interpreted as the extent of contribution each latent variable to phenotype. This can be derived by solving the following variance-covariance model.

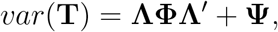

where **Φ** is the variance of latent variables, and **Ψ** is the variance of specific effects (Brown, 2014). Six latent variables were assumed to account for the covariance in the observed phenotypes. Moreover, latent variables were assumed to be correlated with each other. Prior distributions were assigned to all unknown parameters. The non-zero coefficients within factor loading matrix **Λ** were assumed to follow a Gaussian distribution with mean of 0 and variance of 0.01. The variance-covariance matrix **Φ** was assigned an inverse Wishart distribution with a 6 × 6 identity scale matrix **I**_66_ and a degree freedom of 7, 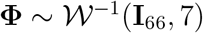 and an inverse Gamma distribution with scale parameter 1 and shape parameter 0.5 was assigned to **Ψ** ~ Γ^−1^(1,0.5).

We employed the blavaan R package (Merkle and Rosseel, 2018) jointly with JAGS (Hornik et al., 2003) to fit the above BCFA. The blavaan runs the runjags R package (Denwood, 2016) to summarize the Markov chain Monte Carlo (MCMC) and samples unknown parameters from the posterior distributions. Three MCMC chains, each of 5,000 samples with 2,000 burn-in, were used to infer the unknown model parameters. The convergence of the parameters was investigated with trace plots and potential scale reduction factor (PSRF) less than 1.2 (Brooks and Gelman, 1998). The PSRF computes the difference between estimated variances among multiple Markov chains and estimated variances within the chain. A large difference indicates non-convergence and may require additional Gibbs sampling.

Subsequently, the posterior means of factor scores (**F**), which reflect the contribution of latent variables to each accession were estimated. Within each draw of Gibbs sampling, **F** was sampled from the conditional distribution of *p*(**F|θ, T**), where ***θ*** refers to the unknown parameters in **Λ, Φ**, and **Ψ**. This conditional distribution was derived with data augmentation (Tanner and Wong, 1987) assuming **F** as missing data (Lee and Song, 2012).

### Multivariate genomic best linear unbiased prediction

We fitted a Bayesian multivariate genomic best linear unbiased prediction to separate breeding values from population structure and noise in the six factor scores computed previously.

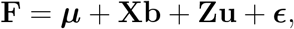

where **μ** is the vector of intercept, **X** is the incidence matrix of covariates, **b** is the vector of covariate effects, **Z** is the incidence matrix relating accessions with additive genetic effects, u is the vector of additive genetic effects, and **ϵ** is the vector of residuals. The incident matrix **X** included subpopulation information (temperate japonica, tropical japonica, indica, aus, aromatic, and admixture), as the rice diversity panel used herein shows a clear substructure (Zhao et al., 2011).

A flat prior was assigned to ***μ*** and **b**, and the joint distribution of **u** and **ϵ** follows multivariate normal

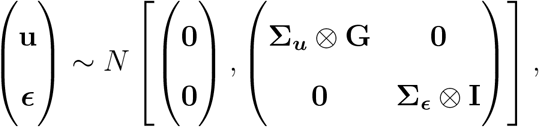

where **G** represents the second genomic relationship matrix of VanRaden (2008), **I** is the identity matrix, Σ_*u*_ and Σ_*ϵ*_ refer to 6 × 6 dimensional genetic and residual variance-covariance matrices, respectively. An inverse Wishart distribution with a 6 × 6 identity scale matrix of **I**_66_ and a degree of freedom 6 was assigned as prior for 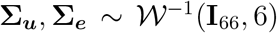. These parameters were selected so that relatively uninformative priors were used. The Bayesian multivariate genomic best linear unbiased prediction model was implemented using the MTM R package (https://github.com/QuantGen/MTM). Posterior mean estimates of genomic correlation between latent variables and predicted breeding values (**û**) were then obtained. The convergence of the estimated parameters was verified by trace plots.

### Sample independence in the Bayesian network

Theoretically, BN learning algorithms assume sample independence. In the multivariate genomic best linear unbiased prediction, the residuals between phenotypes were assumed independent through **I_374×374_**. However, phenotypic dependencies were introduced by the **G** matrix for the additive genetic effects, thereby potentially serving as a confounder. Thus, a transformation of **û** was carried out to derive an adjusted **û**\* by eliminating the dependencies in **G**. For a single trait model, the adjusted **û**\* can be computed by premultiplying **û** by **L**^−1^, where **L** is a lower triangular matrix derived from the Choleskey decompostion of **G** matrix (**G** = **LL**’). Since 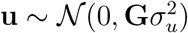, the distribution of **û**\* follows 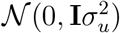 (Callanan and Harville, 1989; Vazquez et al., 2010)

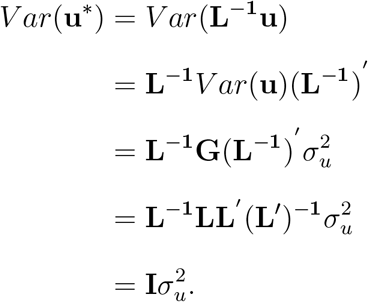

This transformation can be extended to a multi-traits model by defining **u**\* = **M**^−1^**u**, where **M**^−1^ = **I_qq_** ⊗ **L**^−1^ (Töpner et al., 2017). Under the multivariate framework, **u** follows 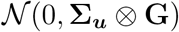 and the variance of **u**\* is

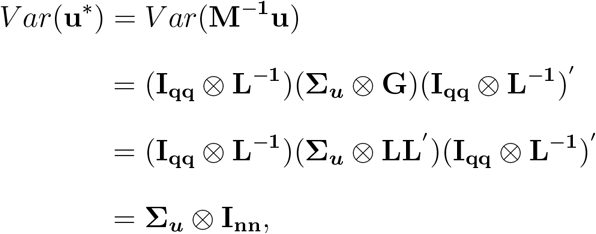

where **L**^−1^**LL**′(**L**^−1^)′ = **I_nn_**. This adjusted **û**\* was used to learn BN structures between predicted breeding values.

### Bayesian network

A BN depicts the joint probabilistic distribution of random variables through their conditional independencies (Scutari and Denis, 2014)

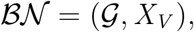

where 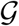 represents a DAG = (*V, E*) with nodes (*V*) connected by one or more edges (*E*) conveying the probabilistic relationships and the random vector *X_V_* = (*X*_1_,…,*X_K_*) is *K* random variables. The joint probability distribution can be factorized as

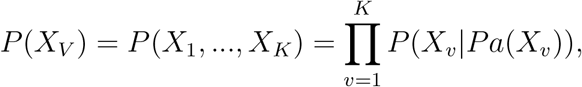

where *Pa*(*X_v_*) denotes a set of parent nodes of child node *X_v_*. The DAG and joint probability distribution are governed by the Markov condition, which states that every random variable is independent of its non-descendants conditioned on its parents. A BN is known as a Gaussian BN, when all variables or phenotypes are defined as marginal or conditional Gaussian distribution as in the present study.

The adjusted breeding values **û**\* were used to infer a genomic network structure among the aforementioned six latent variables. There are three types of structure-learning algorithms for BN: constraint-based algorithms, score-based algorithms, and a hybrid of these two (Scutari and Denis, 2014). The constraint-based algorithms can be originally traced to the inductive causation algorithm (Verma and Pearl, 1991), which uses conditional independence tests for network inference. Briefly, the first step is to identify a d-separation set for each pair of nodes and confer an undirected edge between the two if they are not d-separated. The second step is to identify a v-structure for each pair of non-adjacent nodes, where a common neighbor is the outcome of two non-adjacent nodes. In the last step, compelled edges were identified and oriented, where neither cyclic graph nor new v-structures are permitted. The score-based algorithms are based on heuristic approaches, which first assign a goodness-of-fit score for an initial graph structure and then maximize this score by updating the structure (i.e., add, delete, or reverse the edges of initial graph). The hybrid algorithm includes two steps, restrict and maximize, which harness both constrain-based and score-based algorithms to construct a reliable network. In this study, the two score-based (Hill Climbing and Tabu) and two hybrid algorithms (Max-Min Hill Climbing and General 2-Phase Restricted Maximization) were used to perform structure learning.

We quantified the strength of edges and uncertainty regarding the direction of networks, using 500 bootstrapping replicates with a size equal to the number of accessions and performed structure learning for each replicate in accordance with Scutari and Denis (2014). Non-parametric bootstrap resampling aimed at reducing the impact of the local optimal structures by computing the probability of the arcs and directions. Subsequently, 500 learned structures were averaged with a strength threshold of 85% or higher to produce a more robust network structure. This process, known as model averaging, returns the final network with arcs present in at least 85% among all 500 networks. Candidate networks were compared on the basis of the Bayesian information criterion (BIC) and Bayesian Gaussian equivalent score (BGe). The BIC accounts for the goodness-of-fit and model complexity, and BGe aims at maximizing the posterior probability of networks per the data. All BN were learned via the bnlearn R package (Scutari, 2010). In bnlearn, the BIC score is rescaled by −2, which indicates that the larger BIC refers to a preferred model.

### Data availability

Genotypic data regarding the rice accessions can be downloaded from the rice diversity panel website (http://www.ricediversity.org/). Phenotypic data used herein are available in Zhao et al. (2011), Campbell et al. (2017b), and Supplementary File S3.

## Results

To elucidate the genetic interdependencies among traits typically recorded in breeding programs, we utilized a collection of 48 publicly available phenotypes recorded on a panel of diverse rice accessions (Zhao et al., 2011; Campbell et al., 2017a). The phenotypic data was derived from two independent studies. The first set of phenotypes was recorded from materials grown in two field environments in Arkansas and Faridpur Bangladesh, and in a greenhouse in Aberdeen, UK (Zhao et al., 2011). The 34 phenotypes were recorded at maturity and were largely associated with yield (panicle characteristics flowering time, plant morphology (e.g. height and growth habits), and seed morphological traits. The second study consisted of 14 phenotypes were recorded in a greenhouse environment on plants in the active tillering stage (e.g. 30 day-old plants) under control and saline (14 days of 9.5 dS m−2 NaCl stress). The phenotypes from this study can be classified into three categories: morphological traits (e.g. shoot and root biomass, and plant height), morphological responses to salinity (e.g. the ratio of morphological traits in saline conditions to control), and the ionic components of salinity stress (e.g. Na^+^, K^+^, and Na^+^:K^+^ in both root and shoot tissues) (Campbell et al., 2017a). The complete data set provides an in-depth characterization of phenotypic performance at vegetative and reproductive stages in rice using several classes of traits.

### Latent variable modeling

The BCFA model grouped the observed phenotypes into the underlying latent variables on the basis of prior biological knowledge, assuming these latent variables determine the observed phenotypes. This allowed us to study the genetics of each latent variable. A measurement model derived from BCFA evaluating the six latent variables is shown in Figure 1. Forty-eight observed phenotypes were hypothesized to result from the six latent variables: 7 for flowering time, 14 for morphology, 5 for yield, 11 for grain morphology, 6 for physiology, and 5 for salt response. The convergence of the parameters was confirmed graphically with the trace plots and a PSRF value less than 1.2 (Brooks and Gelman, 1998; Merkle and Rosseel, 2018).

**Figure 1:**
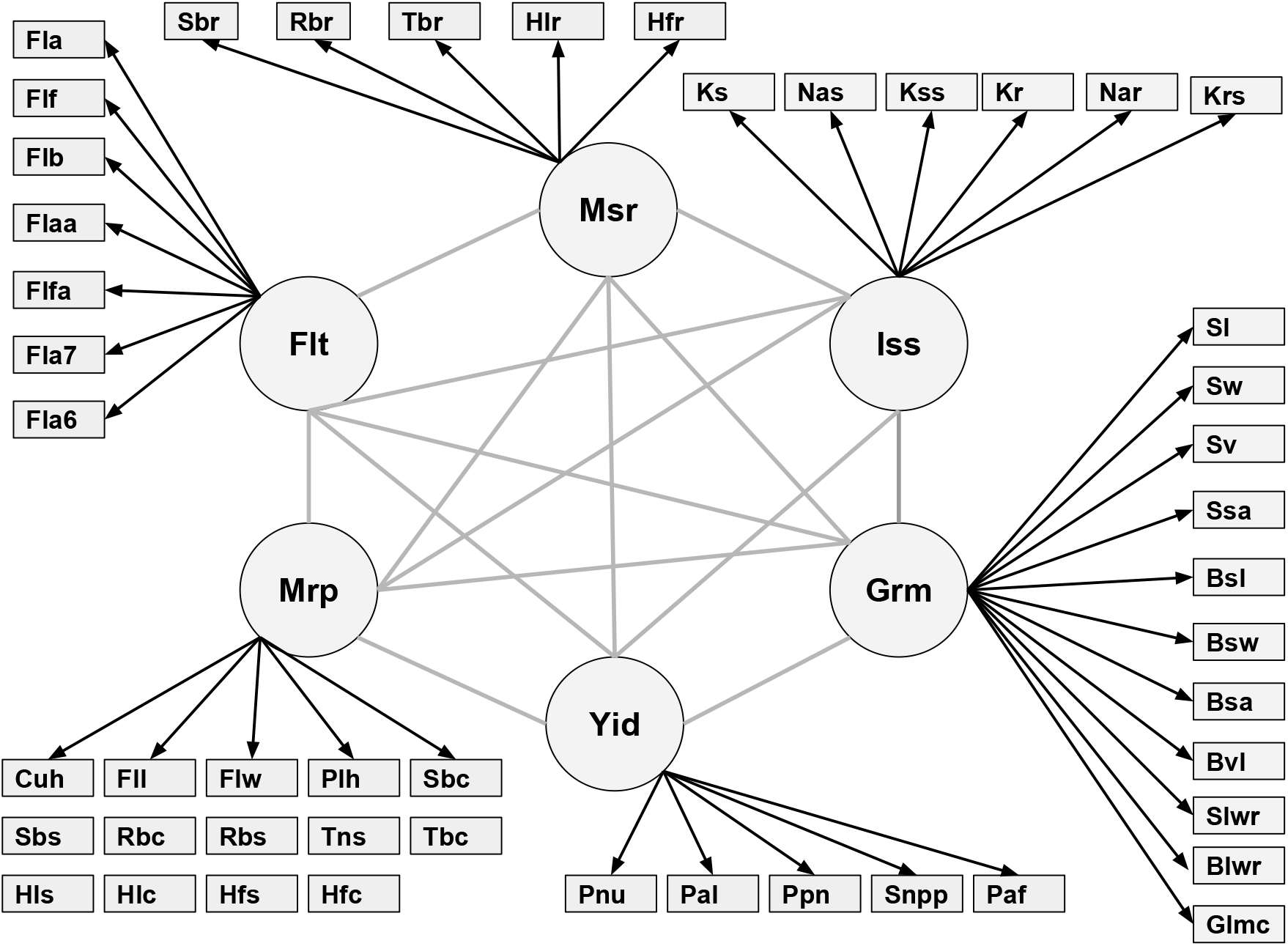
Relationship between six latent variables and observed phenotypes. Msr: morphological salt response; Iss: ionic components of salt stress; Grm: grain morphology; Yid: yield; Mrp: morphology; Flt: flowering time. Abbreviations of observed phenotypes are shown in Table SI.

The six latent factors showed strong contributions to the 48 observed phenotypes, with standardized regression coefficients ranging from −0.668 to 0.980 for flowering time, −0.112 to 0.903 for morphology, −0.113 to 0.977 for yield, −0.501 to 0.986 for grain morphology, −0.016 to 0.829 for physiology, and 0.011 to 0.929 for salt response. The latent factor flowering time showed a strong positive contribution to flowering time in Arkansas (Fla) and Flowering time in Arkansas in 2007 (Fla7) (0.99 and 0.926, respectively; Table 1, indicating that larger values for the latent factor can be interpreted as a greater number of days from sowing to emergence of the inflorescence. The latent factor morphology showed the largest positive contributions to traits describing height during the vegetative stage (e.g. height to newest ligule in salt (Hls), 0.920; height to newest ligule in control (Hlc), 0.899; height to the tip of first fully expanded leaf in salt (Hfs), 0.907; and height to tip of first fully expanded leaf in control (Hfc)), 0.925; suggesting that this latent factor is an overall representation of plant size. Yield showed large positive contributions to the observed phenotypes primary panicle branch number (Ppn) and seed number per panicle (Snpp) (0.790 and 0.780, respectively), suggesting that larger values for yield indicate a higher degree of branching and seed number. Observed phenotypes describing seed size (e.g. seed volume (Sv) and brown rice volume (Bvl) (0.990 and 0.986, respectively)) were most strongly associated with grain morphology. The latent factor ionic components of salt stress showed strong positive contributions to two observed phenotypes that quantify the ionic components of salt stress (shoot Na^+^:K^+^ (Kslm) and shoot Na^+^ (Nas) (0.983 and 0.975, respectively), indicating that higher values for the latent factor result in greater shoot Na^+^ and Na^+^:K^+^. Finally, the latent factor describing morphological salt response showed strong positive contributions to the observed phenotype describing the effect of salt treatment on plant height (ratio of height to tip of newest fully expanded leaf in salt to that of control plants (Hfr) (0.939), thus larger values for the latent factor may indicate a more tolerant growth response to salinity.

**Table 1:**
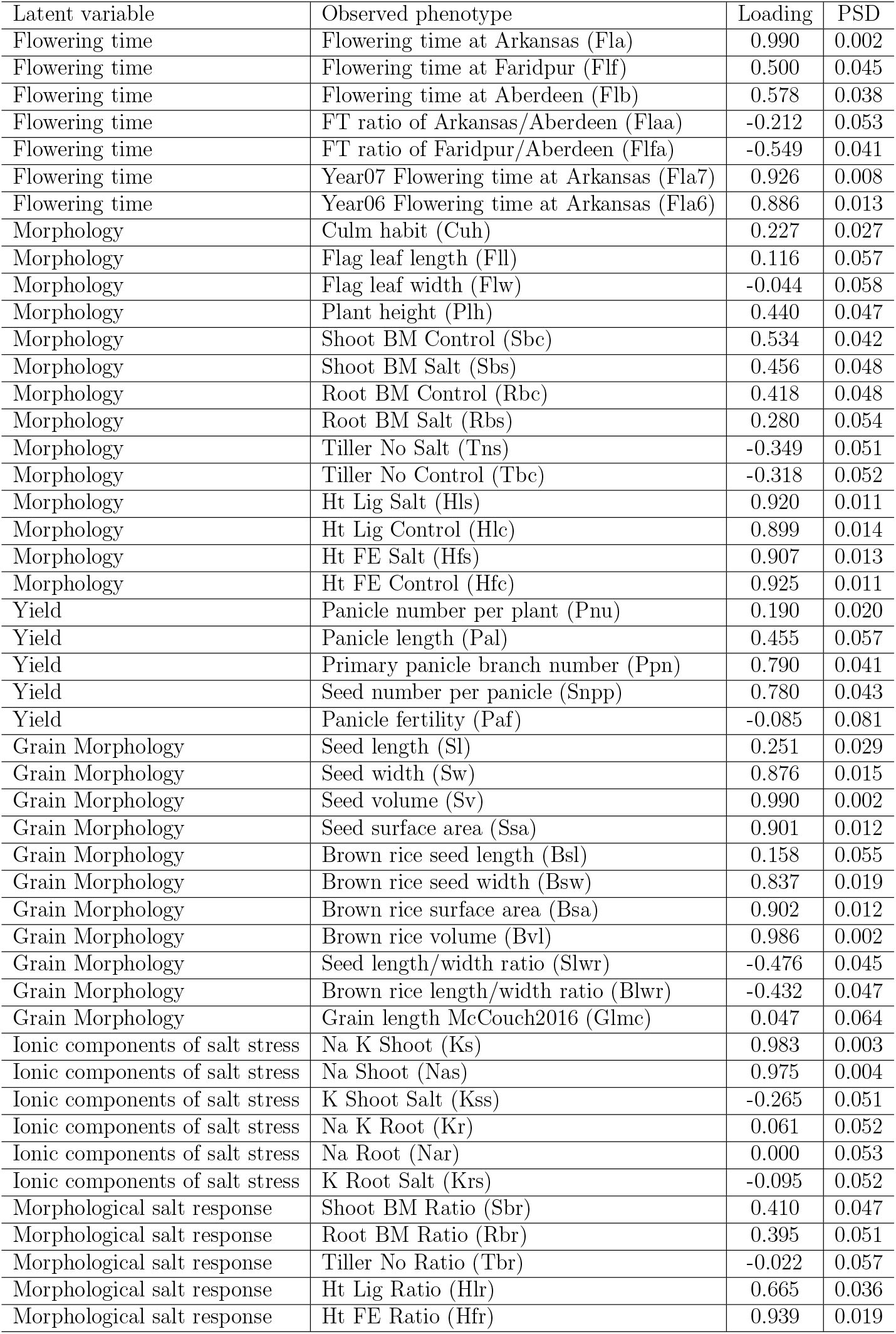
Standardized factor loadings obtained from the Bayesian confirmatory factor analysis. PSD refers to the posterior standard deviation of standardized factor loadings.

### Genomic correlation among latent variables

To understand the genetic relationships between latent variables, genomic correlation analysis was performed. Genomic correlation is due to pleiotropy or linkage disequilibrium between QTL. The genomic correlations among latent variables are shown in Figure 2. Negative correlations were observed between salt response (Slr) and all other five latent variables. In particular, flowering time (−0.5), yield (−0.54), and grain morphology (−0.74) were negatively correlated with morphological salt response 2. These results suggest that accessions that harbor alleles for more tolerant morphological salt responses may also have alleles associated with longer flowering times, smaller seeds, and low yield. Similarly, a negative correlation was observed between morphology and yield (−0.56) and between morphology and grain morphology (−0.31). Thus, accessions with alleles associated with large plant size may also have alleles that result in low yield, small grain volume, and lower shoot Na^+^ and Na^+^:K^+^. In contrast, a positive correlation was observed between grain morphology and yield (0.49) and between grain morphology and ionic components of salt stress (0.4). Thus, selection for large grain may result in improved yield, and higher shoot Na^+^ and Na^+^:K^+^.

**Figure 2:**
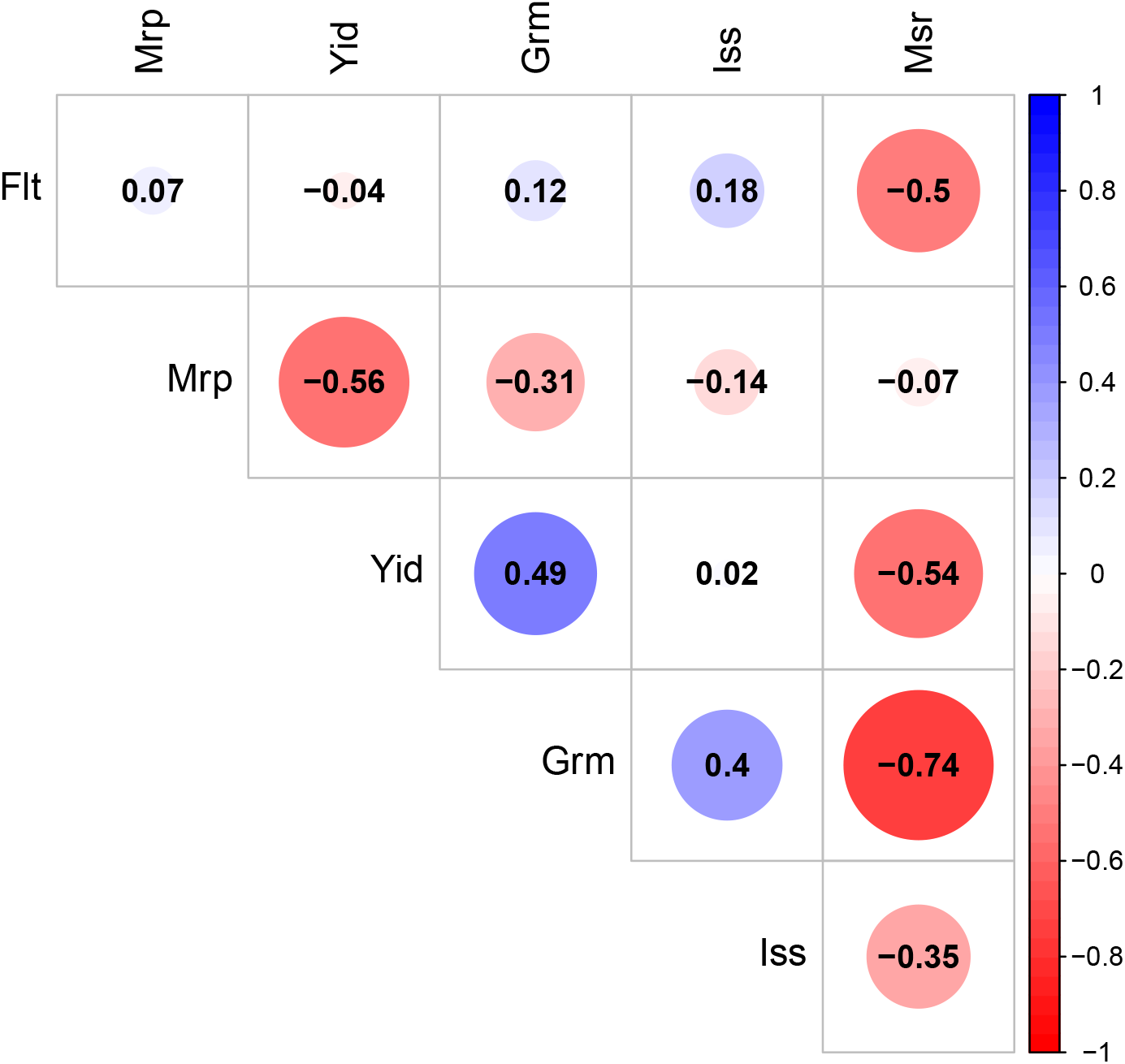
Genomic correlation of six latent variables. The size of each circle, degree of shading, and value reported correspond to the correlation between each pair of latent variables. Msr: morphological salt response; Iss: ionic components of salt stress; Grm: grain morphology; Yid: yield; Mrp: morphology; Flt: flowering time.

### Bayesian network

To infer the possible causal structure between latent variables, BN was performed. Prior to BN, the normality of latent variables was assessed using histogram plots combined with density curves as shown in Figure S2. Overall, all the six latent variables approximately followed a Gaussian distribution.

The Bayesian networks learned with the score-based and hybrid algorithms are shown in Figures 3, 4, 5, and 6. The structures of BN were refined by model averaging with 500 networks from bootstrap resampling to reduce the impact of local optimal structures. The labels of the arcs measure the uncertainty of the arcs, corresponding to strength and direction (in parenthesis). The former measures the frequency of the arc presented among all 500 networks from the bootstrapping replicates and the latter is the frequency of the direction shown conditional on the presence of the arc. We observed minor differences in the structures presented within and across the two types of algorithms used. In general, small differences were observed within algorithm types compared to those across algorithms. The two score-based algorithms produced a greater number of edges than two hybrid algorithms. In Figure 3, the Hill Climbing algorithm produced seven directed connections among the six latent variables. Three connections were indicated towards flowering time from morphological salt response, ionic components of salt stress, and morphology, and two edges to yield from morphology and from grain morphology. Other two edges were observed from ionic components of salt stress to grain morphology and from grain morphology to morphological salt response. A similar structure was generated by the Tabu algorithm, except that the connection between salt response and grain morphology presented an opposite direction (Figure 4). The Max-Min Hill Climbing hybrid algorithm yielded six directed edges from morphological salt response to grain morphology, from ionic components of salt stress to grain morphology, from ionic components of salt stress to flowering time, from flowering time to morphology, from morphology to yield, and from grain morphology to yield (Figure 5). An analogous structure with the only difference observed in the directed edge from morphology to flowering time was inferred with the General 2-Phase Restricted Maximization algorithm as shown in Figure 6. Across all four algorithms, there were four common directed edges: from ionic components of salt stress to flowering time and to grain morphology, and from morphology and grain morphology to yield. The most favorable network was considered the one from the Tabu algorithm, which returned the largest network score in terms of BIC (1086.61) and BGe (1080.88). Collectively, these results suggest that there may be a direct genetic influence of morphology and grain morphology on yield, and physiological components of salt tolerance on grain morphology and flowering time.

**Figure 3:**
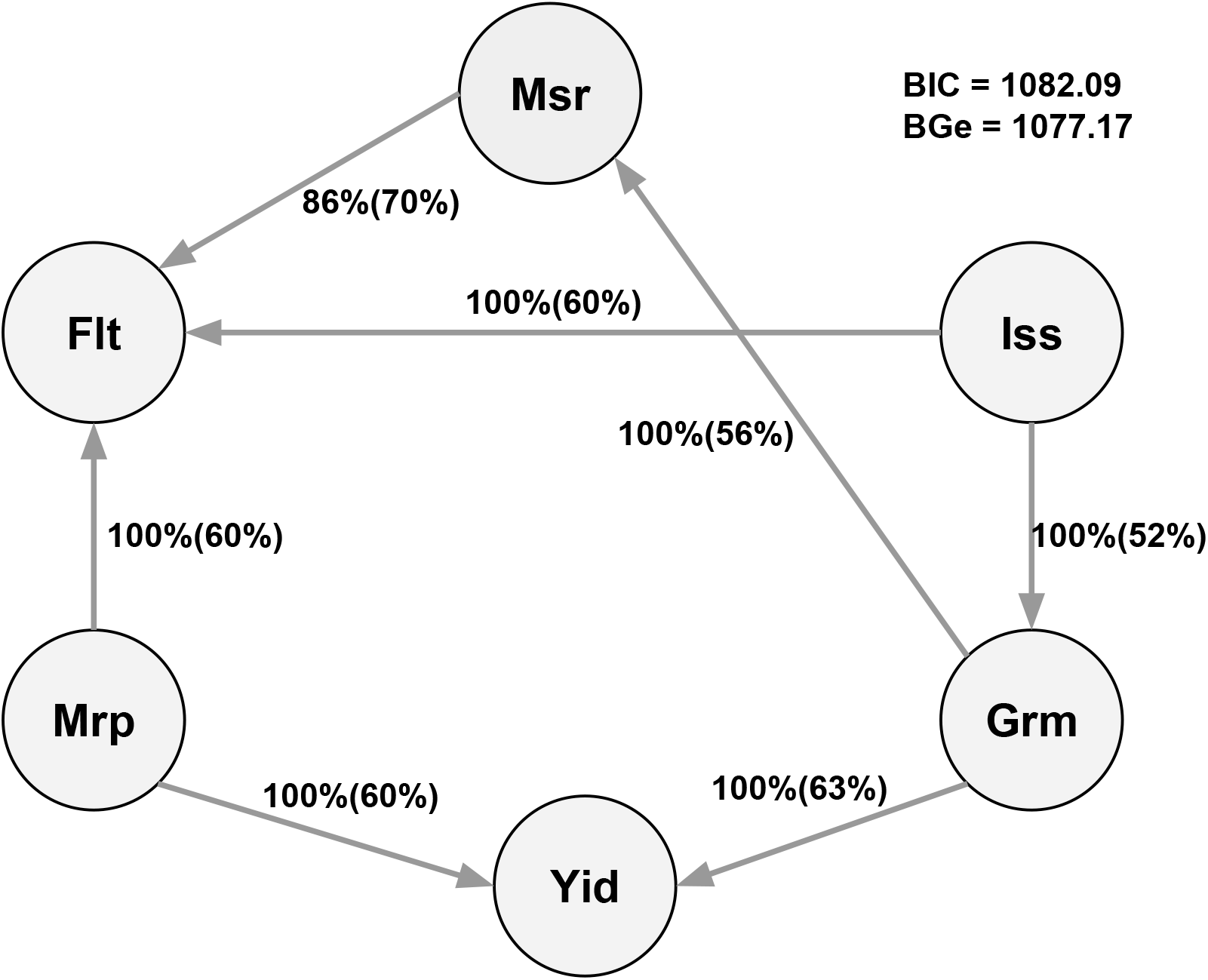
Bayesian network between six latent variables based on the Hill Climbing algorithm. The quality of the structure was evaluated by bootstrap resampling and model averaging across 500 replications. Labels of the edges refer to the strength and direction (parenthesis) which measure the confidence of the directed edge. The strength indicates the frequency of the edge is present and the direction measures the frequency of the direction conditioned on the presence of edge. BIC: Bayesian information criterion score. BGe: Bayesian Gaussian equivalent score. Msr: morphological salt response; Iss: ionic components of salt stress; Grm: grain morphology; Yid: yield; Mrp: morphology; Flt: flowering time.

**Figure 4:**
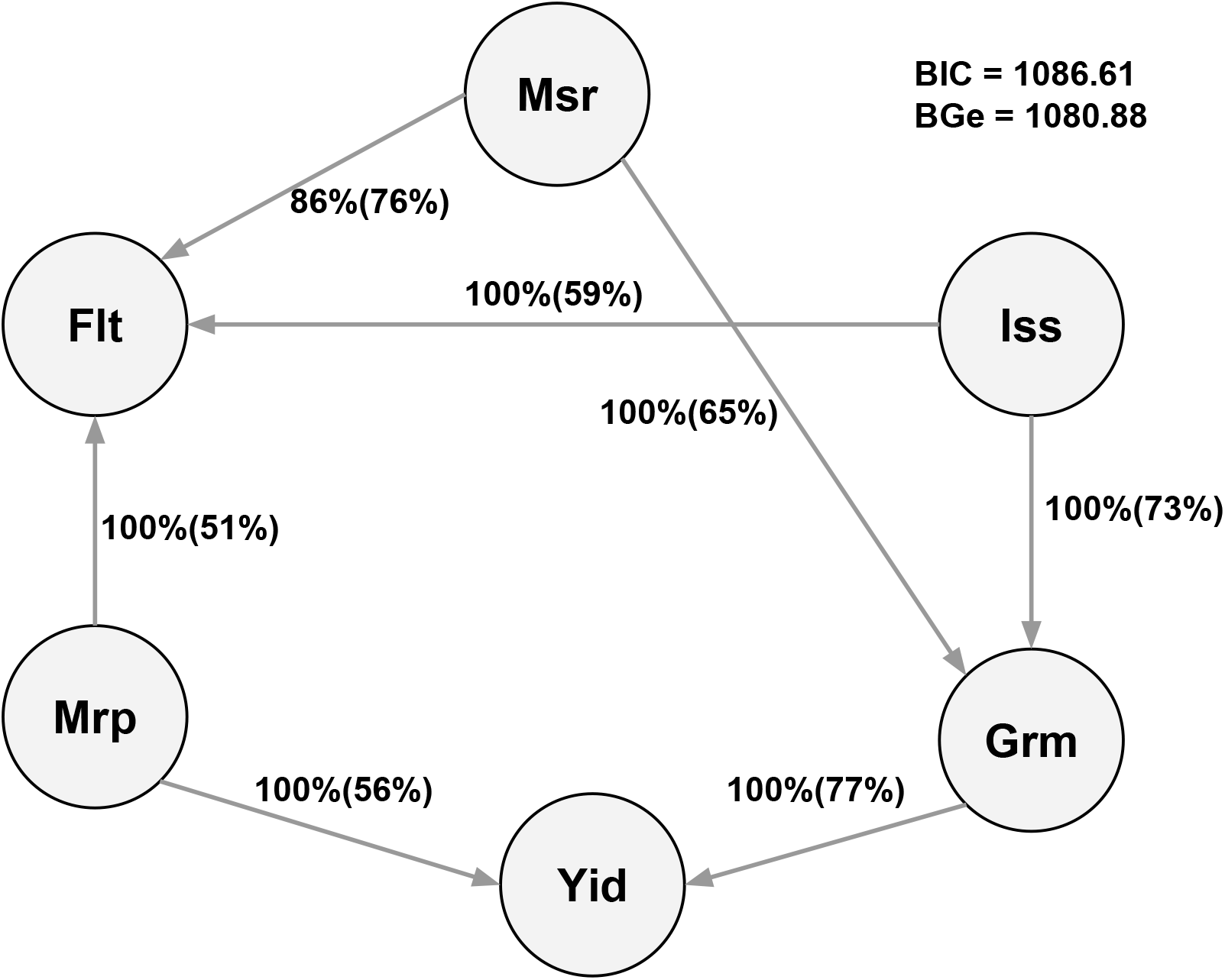
Bayesian network between six latent variables based on the Tabu algorithm. The quality of the structure was evaluated by bootstrap resampling and model averaging across 500 replications. Labels of the edges refer to the strength and direction (parenthesis) which measure the confidence of the directed edge. The strength indicates the frequency of the edge is present and the direction measures the frequency of the direction conditioned on the presence of edge. BIC: Bayesian information criterion score. BGe: Bayesian Gaussian equivalent score. Msr: morphological salt response; Iss: ionic components of salt stress; Grm: grain morphology; Yid: yield; Mrp: morphology; Flt: flowering time.

**Figure 5:**
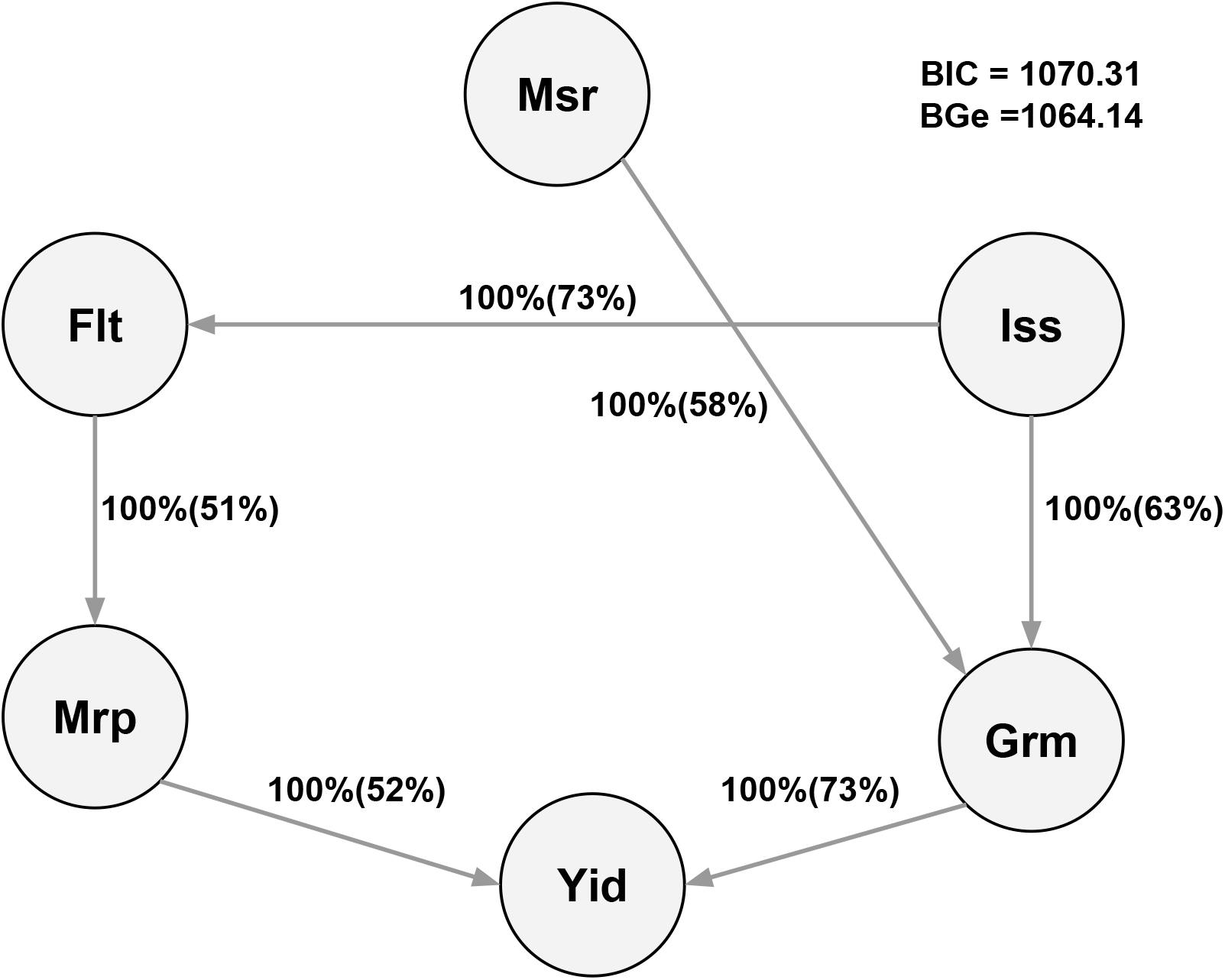
Bayesian network between six latent variables based on the Max-Min Hill Climbing algorithm. The quality of the structure was evaluated by bootstrap resampling and model averaging across 500 replications. Labels of the edges refer to the strength and direction (parenthesis) which measure the confidence of the directed edge. The strength indicates the frequency of the edge is present and the direction measures the frequency of the direction conditioned on the presence of edge. BIC: Bayesian information criterion score. BGe: Bayesian Gaussian equivalent score. Msr: morphological salt response; Iss: ionic components of salt stress; Grm: grain morphology; Yid: yield; Mrp: morphology; Flt: flowering time.

**Figure 6:**
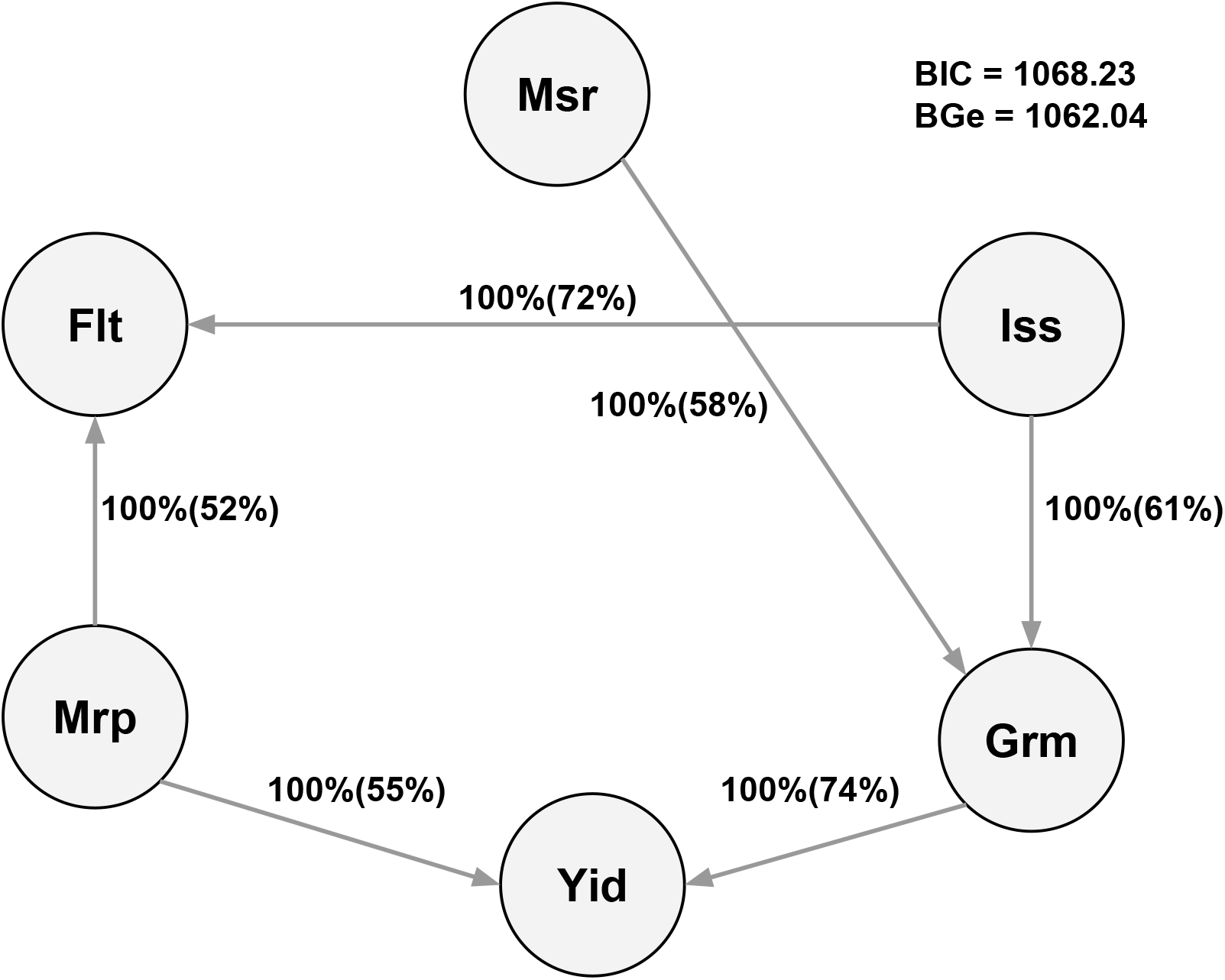
Bayesian network between six latent variables based on the General 2-Phase Restricted Maximization algorithm. The quality of the structure was evaluated by bootstrap resampling and model averaging across 500 replications. Labels of the edges refer to the strength and direction (parenthesis) which measure the confidence of the directed edge. The strength indicates the frequency of the edge is present and the direction measures the frequency of the direction conditioned on the presence of edge. BIC: Bayesian information criterion score. BGe: Bayesian Gaussian equivalent score. Msr: morphological salt response; Iss: ionic components of salt stress; Grm: grain morphology; Yid: yield; Mrp: morphology; Flt: flowering time.

## Discussion

This study is based on the premise that most phenotypes interact to greater or lesser degrees with each other through underlying physiological and molecular pathways. While these physiological pathways are important for the development of agronomically important characteristics, they are often unknown or difficult to assess in large populations. The approach utilized here leverages phenotypes that can be readily assessed in large populations to quantify these underlying unobserved phenotypes, and elucidates the relationships between these variables.

Understanding the behaviors among phenotypes in the complex traits is critical for genetic improvement of agricultural species (Hickey et al., 2017). Graphical modeling offers an avenue to decipher bi-directional associations or probabilistic dependencies among variables of interest in plant and animal breeding. For instance, BN and L1-regularized undirected network can be used to model interrelationships of linkage disequilibrium (LD) (Morota et al., 2012; Morota and Gianola, 2013) or phenotypic, genetic, and environmental interactions (Xavier et al., 2017) in a systematic manner. Importantly, MTM elucidates both direct and indirect relationships among phenotypes. Inaccurate interpretation of these relationships may substantially bias selection decisions (Valente et al., 2015; Gianola et al., 2015). Thus, we applied BCFA to reduce the dimension of the responses by hypothesizing 48 manifest phenotypes originated from the underlying six constructed latent variables as shown in Figure 1 assuming that these latent traits are most important, followed by application of BN to infer the structures among the six biologically relevant latent variables (Figures 3,4, 5, and 6). The BN represents the conditional dependencies between variables. Care must be taken in interpreting these relationships as a causal effect. Although a good BN is expected to describe the underlying causal structure per the data, when the structure is learned solely on the basis of the observed data, it may return multiple equivalent networks that describe the data well. In practice, searching such a causal structure with observed data needs three additional assumptions (Scutari and Denis, 2014): 1) each variable is independent of its non-effects (i.e., direct and indirect) conditioned on its direct causes, 2) the probability distribution of variables is supported by a DAG, where the d-separation in DAG provides all dependencies in the probability distribution, and 3) no additional variables influence the variables within the network. Although it may be difficult to meet these assumptions in the observed data, a BN is equipped with suggesting potential causal relationships among latent variables, which can assist in exploring data, making breeding decisions, and improving management strategies in breeding programs (Rosa et al., 2011).

### Biological meaning of latent variables and their relationships

We performed BCFA to summarize the original 48 phenotypes with the six latent variables. The number of latent variables and which latent variables load onto phenotypes were determined from the literature. The latent variable morphological salt response (Slr) contributed strongly to salt indices for shoot biomass, root biomass, and two indices for plant height (Table 1). Thus, morphological salt response can be interpreted as the morphological responses to salinity stress, with higher values indicating a more tolerant growth response. The latent variable yield is a representation of overall grain productivity, and contributed strongly to the observed phenotypes primary panicle branch number, seed number per panicle, and panicle length. The positive loading scores on these observable phenotypes indicates that more highly branched, productive panicles will have higher values for yield (Table 1). Seed width, seed volume, and seed surface area contributed significantly to the latent variable grain morphology (Grm) (Table 1). Therefore, these results indicate that the grain morphology is a summary of the overall shape of the grain, where high values represent large, round grains, while low values represent small, slender grains. Considering the grain characteristics of rice subpopulations, temperate japonica accessions are expected to have high values for grain morphology, while indica accessions have lower values for grain morphology. Latent variable morphology (Mrp) is a representation of plant biomass during the vegetative stage (28-day-old plants) (Table 1). Shoot biomass, root biomass, and two metrics for plant height contributed largely to morphology, suggesting that accessions with high values for morphology are tall plants with a large biomass.

Genomic correlation analysis among the six latent variables showed meaningful correlations among several pairs. These genetic correlations can either be caused by linkage or pleiotropy. The former is likely to prevail in species with high LD, which is the case in rice where LD ranges from 100 to 200kb (Huang et al., 2010). A negative relationship was observed between morphological salt response and three other latent variables (Figure 2). For instance, a negative correlation between morphological salt response and yield indicates that accessions of samples harboring alleles for superior morphological salt responses (e.g. those that are more tolerant) tend to also harbor alleles for poor yield (Figure 2). The rice diversity panel we used is a representative sample of the total genetic diversity within cultivated rice and contains many unimproved traditional varieties (~12% of lines in the study are landraces and ~33% classified as cultivars; Supplementary File S2) and modern breeding lines (Eizenga et al., 2014). While traditional varieties exhibit superior adaptation to abiotic stresses, they often have very poor agronomic characteristics including low yield, late flowering, and high photoperiod sensitivity (Thomson et al., 2009, 2010). Moreover, the indica and japonica subspecies have contrasting salt responses and very different grain morphology. Japonica accessions tend to have short, round seeds and are more sensitive to salt stress, while indica accessions have long, slender grains and often are more salt tolerant (Zhao et al., 2011; Campbell et al., 2017a). The negative relationship observed between salt response and grain morphology suggests that lines that harbor alleles for high grain morphology (e.g., large, round grains) tend to also harbor alleles for a tolerant growth response to salt stress. However, no studies have yet reported an association between alleles for grain morphology and morphological salt response. Therefore, it remains to be addressed whether this relationship is due to LD or pleitropy.

Genetic correlations observed between other latent variables may suggest a pleiotropic effect among loci. For instance, a negative relationship was observed between morphological salt response and ionic components of salt stress, indicating that accessions harboring alleles associated with superior morphological salt response also tend to harbor alleles for reduced ion content under salt stress (Figure 2). The relationship between salt tolerance, measured in terms of growth or yield, and Na^+^ and Na^+^:K^+^ has been a documented for decades (reviewed by Munns and Tester (2008)). Moreover, natural variation for Na^+^ transporters has been utilized to improve growth and yield under saline conditions in rice and other cereals (Ren et al., 2005; Byrt et al., 2007; Horie et al., 2009; Munns et al., 2012; Campbell et al., 2017a). Therefore, the negative genetic relationships observed between morphological salt response and ion content may be due to the pleiotropic effects of some loci.

The genomic relationships among latent variables including morphology, yield, and grain morphology may have resulted from the selection of alleles associated with good agronomic characteristics. A positive relationship was observed between yield and grain morphology, suggesting that alleles that positively contribute to productive panicles also may contribute to large, round grains. Furthermore, the negative genomic correlation observed between morphology and yield indicates that alleles negatively influencing total plant biomass also have a positive contribution to traits for productive panicles. This genomic relationship may reflect the genetics of harvest index, which is defined as the ratio of grain yield to total biomass. Over the past 50 years, rice breeders have selected high harvest index, resulting in plants with short compact morphology and many highly productive panicles (Hay, 1995; Peng et al., 2008).

Although BCFA may yield biologically meaningful results, a potential limitation of BCFA is that we assumed each phenotype does not measure more than one latent variable. This assumption may not always strictly concur with the observational data. Therefore, further studies are required to allow each phenotype to potentially load onto multiple factors in the BCFA framework. An alternative approach is to derive the number of latent variables and determine which latent variables load onto phenotypes directly from observed data, using exploratory FA. This approach was not pursued here because accurate estimation of unknown parameters in the exploratory FA requires a large sample size, which was not the case herein (Brown, 2014).

### Bayesian network of latent variables

The BN is a probabilistic DAG, which represents the conditional dependencies among phenotypes. The genomic correlation among latent variables described in Figure 2 does not inform the flow of genetic signals nor distinguish direct and indirect associations, whereas BN displays directions between latent variables and separate direct and indirect associations. Therefore, the BN describes the possibility that other phenotypes will change if one phenotype is intervened (i.e., selection). However, caution is required to interpret this network as a causal effect, as the causal BN requires more assumptions, which are usually difficult to meet in observational data (Pearl, 2009).

Four common edges or consensus subnetworks across the four BN may be the most reliable substructure of latent variables and may describe the dependence between agronomic traits (Figures 3, 4, 5, and 6). For example, edges from grain morphology to yield and morphology to yield can be interpreted as final grain productivity is dependant on specific vegetative characteristics as well grain traits. This is because yield, which represents the overall grain productivity of a plant, depends on morphological characteristics such as the degree of tillering, an architecture that allows the plant to efficiently capture light and carbon, and a stature that is resistant to lodging, the degree of panicle branching, as well as specific grain characteristics such as seed volume and shape. Moreover, there is a direct biological linkage between specific vegetative architectural traits such as tillering and plant height, and yield related traits such as panicle branching and number of seeds per panicle. The degree of branching during both vegetative and reproductive development is dependant on the development and initiation of auxiliary meristems. Several genes have been identified in this pathway and have shown to have pleiotropic effects on tillering and panicle branching (reviewed by Liang et al. (2014)). For instance, *OsSPL14* has been shown to be an important regulator of auxiliary branching in both vegetative and reproductive stages in rice (Jiao et al., 2010; Miura et al., 2010). Moreover, other genes such as *OsGhd8* have been reported to regulate other morphological traits such as plant height and yield through increase panicle branching (Yan et al., 2011). The biological importance of these dependencies can also be illustrated by viewing them in the context of genetic improvement, as selection for specific architectural traits (represented by the latent variable morphology) and grain characteristics have traditionally been used as traits to improve rice productivity in many conventional breeding programs (Redona and Mackill, 1998; Huang et al., 2013).

While the above example provides a plausible network structure between latent variables, edges from ionic components of salt stress to flowering time and to grain morphology are an example of instances where caution should be used to infer causation. As mentioned above, there is an inherent difference in salt tolerance and grain morphological traits between the indica and japonica subspecies. The edges observed for these two latent variables (ionic components of salt stress and grain morphology) in BN may be driven by LD between alleles associated with grain morphology and alleles for salt tolerance rather than pleitropy. Thus, given the current data set, genetic effects for grain morphology may still be conditionally dependant on ionic components of salt stress and the BN may be true, even if there is no direct overlap in the genetic mechanisms for the two traits.

We found that there are some uncertain edges among BN. For instance, direction from salt response to grain morphology is supported by 65% (Figure 4), 58% (Figure 5), and 58% (Figure 6) bootstrap sampling, whereas the opposite direction is supported by 56% bootstrap sampling (Figure 3). An analogous uncertainty was also observed between morphology and flowering time, i.e., the path from morphology to flowering time was supported 60% (Figure 3), 51% (Figure 4), and 52% (Figure 6), while the reverse direction was supported 51% (Figure 6) upon bootstrapping. In addition, the two score-based algorithms captured edges between morphological salt response and flowering time with 70% and 76% bootstrapping evidence. However, this connection was not detected in the two hybrid algorithms. In general, inferring the direction of edges was harder than inferring the presence or absence of undirected edges. Finally, the whole structures of BN were evaluated in terms of the BIC score and BGe. Ranking of the networks was consistent across BIC and BGe and the two score-based algorithms produced networks with greater goodness-of-fit than the two hybrid algorithms. The optimal network was produced by the Tabu algorithm. This is consistent with the previous study reporting that the score-based algorithm produced a better fit of networks in data on maize (Topner et al., 2017).

In conclusion, the present results show the utility of CFA and network analysis to characterize various phenotypes in rice. We showed that the joint use of BCFA and BN can be applied to predict the potential influence of external interventions or selection associated with target traits such as yield in the high-dimensional interrelated complex traits system. We contend that the approaches used herein provide greater insights than pairwise-association measures of multiple phenotypes and can be used to analyze the massive amount of diverse image-based phenomics dataset being generated by the automated plant phenomics platforms (e.g., Furbank and Tester, 2011). With a large volume of complex traits being collected through phenomics, numerous opportunities to forge new research directions are generated by using network analysis for the growing number of phenotypes.

