## Supplementary File 1 for "Genomic Bayesian confirmatory factor analysis and Bayesian network to characterize a wide spectrum of rice phenotypes"

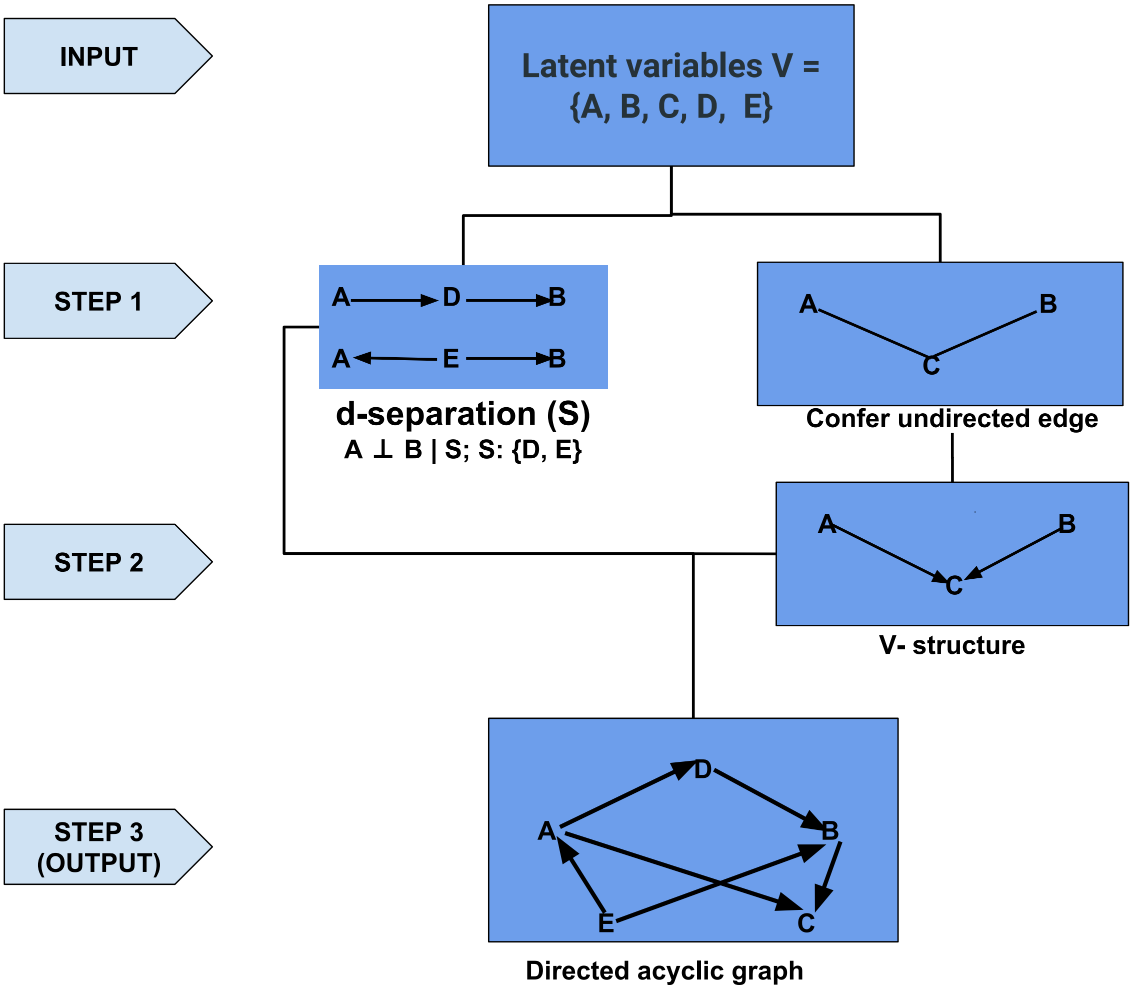


Figure S1: Flow diagram to illustrate the concept of constraint-based structure learning algorithm for a Bayesian network. The A, B, C, D, and E represent five nodes or latent variables. S refers to a set of d-separation.

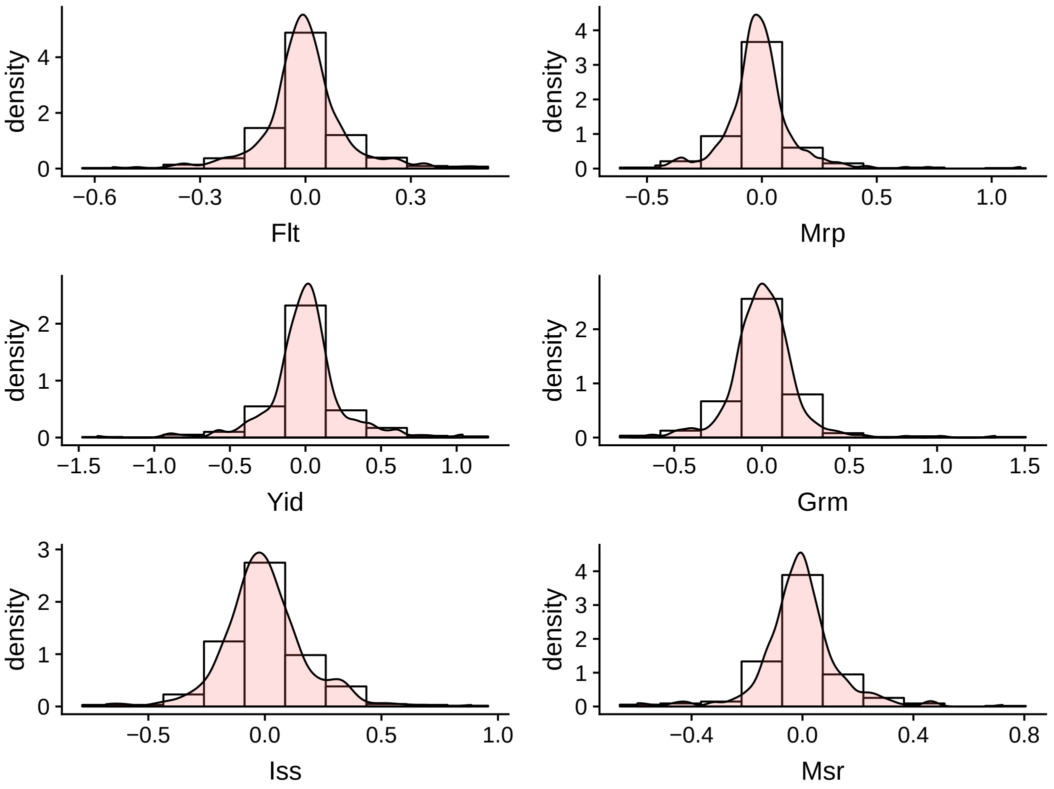


Figure S2: Histogram plots and density curves of six latent variables. Flt: flowering time; Mrp: morphology; Yid: yield; Grm: grain morphology; Iss: ionic components of salt stress; Msr: morphological salt response.

### Table S1: Description of phenotypes used.

| Phenotype Label | Category | Description | Publication |
|---|---|---|---|
| Flowering time at Arkansas (Fla) | Flowering time | Number of days until the inflorescence is 50% emerged from the flag leaf counted from the day of planting. Measured in Arkansas, USA. | Zhao et al. (2011) |
| Flowering time at Faridpur (Flf) | Flowering time | Number of days until the inflorescence is 50% emerged from the flag leaf counted from the day of transplanting. Measured in Faridpur, BD. | Zhao et al. (2011) |
| Flowering time at Aberdeen (Flb) | Flowering time | Number of days until the inflorescence is 50% emerged from the flag leaf counted from the day of planting. Measured in Aberdeen, UK. | Zhao et al. (2011) |
| FT ratio of Arkansas/Aberdeen (Flaa) | Flowering time | Ratio of days to heading in Arkansas to days to heading in Aberdeen. Metric used to describe photoperiod sensitivity. | Zhao et al. (2011) |
| FT ratio of Faridpur/Aberdeen (Flfa) | Flowering time | Ratio of days to heading in Faridpur to days to heading in Aberdeen. Metric used to describe photoperiod sensitivity. | Zhao et al. (2011) |
| Year07 Flowering time at Arkansas (Fla7) | Flowering time | Number of days until the inflorescence is 50% emerged from the flag leaf counted from the day of planting. Measured in Arkansas in 2007. | Zhao et al. (2011) |
| Year06 Flowering time at Arkansas (Fla6) | Flowering time | Number of days until the inflorescence is 50% emerged from the flag leaf counted from the day of planting. Measured in Arkansas in 2006. | Zhao et al. (2011) |
| Seed length (Sl) | Grain morphology | Length of the seed with hull (palea and lemma) | Zhao et al. (2011) |
| Seed width (Sw) | Grain morphology | Width of the seed with hull (palea and lemma) | Zhao et al. (2011) |
| Seed volume (Sv) | Grain morphology | Volume of the seed with hull (palea and lemma) | Zhao et al. (2011) |
| Seed surface area (Ssa) | Grain morphology | Surface area of the seed with hull (palea and lemma) | Zhao et al. (2011) |
| Brown rice seed length (Bsl) | Grain morphology | Length of the unpolished rice grain (dehulled seed) | Zhao et al. (2011) |
| Brown rice seed width (Bsw) | Grain morphology | Width of the unpolished rice grain (dehulled seed) | Zhao et al. (2011) |
| Brown rice surface area (Bsa) | Grain morphology | Surface area of the unpolished rice grain (dehulled seed) | Zhao et al. (2011) |
| Brown rice volume (Bvl) | Grain morphology | Volume of the unpolished rice grain (dehulled seed) | Zhao et al. (2011) |
| Seed length/width ratio (Slwr) | Grain morphology | Ratio of seed length/ seed width (with hull) | Zhao et al. (2011) |
| Brown rice length/width ratio (Blwr) | Grain morphology | Ratio of unpolished rice grain length/grain width (dehulled seed) | Zhao et al. (2011) |
| Grain length McCouch2016 (Glmc) | Grain morphology | Length of the seed with hull (palea and lemma). Reported by McCouch et al (JSJSJ) | McCouch et al. (2016) |
| Culm habit (Cuh) | Morphology | Average culm angle of plants at maturity | Zhao et al. (2011) |
| Flag leaf length (Fll) | Morphology | Length of the flag leaf measured from leaf base to leaf tip (cm) | Zhao et al. (2011) |
| Flag leaf width (Flw) | Morphology | Width of the flag leaf measured at the widest portion of flag leaf lamina (cm) | Zhao et al. (2011) |
| Plant height (Plh) | Morphology | Height of plant from soil surface to tip of main panicle (inflorescence) (cm) | Zhao et al. (2011) |
| Shoot BM Control (Sbc) | Morphology | Shoot dry weight (g) for 28 day old plants in control. | Campbell et al. (2017) |
| Shoot BM Salt (Sbs) | Morphology | Shoot dry weight (g) in for 28 day old plants 9 dS salt. | Campbell et al. (2017) |
| Root BM Control (Rbc) | Morphology | Root dry weight (g) for 28 day old plants in control. | Campbell et al. (2017) |
| Root BM Salt (Rbsl) | Morphology | Root dry weight (g) for 28 day old plants in 9 dS salt. | Campbell et al. (2017) |
| Tiller No Salt (Tns) | Morphology | Tiller number for 28 day old plants in 9 dS salt. | Unpublished. Study described in Campbell et al. (2017) |
| Tiller No Control (Tbc) | Morphology | Tiller number for 28 day old plants in control. | Unpublished. Study described in Campbell et al. (2017) |
| Ht Lig Salt (Hls) | Morphology | Height (cm) from the soil to the ligule of the newest expanded leaf in salt. Measured on 28 day old plants. | Unpublished. Study described in Campbell et al. (2017) |
| Ht Lig Control (Hlc) | Morphology | Height (cm) from the soil to the ligule of the newest expanded leaf in salt. Measured on 28 day old plants. | Unpublished. Study described in Campbell et al. (2017) |
| Ht FE Salt (Hfs) | Morphology | Height (cm) from the soil to the tip of the newest expanded leaf in salt. | Unpublished. Study described in Campbell et al. (2017) |
| Ht FE Control (Hfc) | Morphology | Height (cm) from the soil to the tip of the newest expanded leaf in control. Measured on 28 day old plants. | Unpublished. Study described in Campbell et al. (2017) |
| Na K Shoot (Ks) | Ionic components of salt stress | Shoot sodium to potassium ratio ($10^3$) in salt. Measured on 28 day old plants after 14 days of 9dS salt stress. | Campbell et al. (2017) |
| Na Shoot (Nas) | Ionic components of salt stress | Shoot sodium content (mmol/g dry wt.) in salt. Measured on 28 day old plants after 14 days of 9dS salt stress. | Campbell et al. (2017) |
| K Shoot Salt (Kss) | Ionic components of salt stress | Shoot potassium content (mmol/g dry wt.) in salt. Measured on 28 day old plants after 14 days of 9dS salt stress. | Campbell et al. (2017) |
| Na K Root (Kr) | Ionic components of salt stress | Root sodium to potassium ratio ($10^3$) in salt. Measured on 28 day old plants after 14 days of 9dS salt stress. | Campbell et al. (2017) |
| Na Root (Narl) | Ionic components of salt stress | Root sodium content (mmol/g dry wt.) in salt. Measured on 28 day old plants after 14 days of 9dS salt stress. | Campbell et al. (2017) |
| K Root Salt (Krsl) | Ionic components of salt stress | Root potassium content (mmol/g dry wt.) in salt. Measured on 28 day old plants after 14 days of 9dS salt stress. | Campbell et al. (2017) |
| Shoot BM Ratio (Sbr) | Morphological salt response | Ratio of the LSMeans for shoot dry in salt over control. Measured on 28 day old plants after 14 days of 9dS salt stress. | Campbell et al. (2017) |
| Root BM Ratio (Rbr) | Morphological salt response | Ratio of the LSMeans for root dry in salt over control. Measured on 28 day old plants after 14 days of 9dS salt stress. | Campbell et al. (2017) |
| Tiller No Ratio (Tbr) | Morphological salt response | Ratio of the tiller number in salt over tiller number in control. Measured on 28 day old plants after 14 days of 9dS salt stress. | Unpublished. Study described in Campbell et al. (2017) |
| Ht Lig Ratio (Hlr) | Morphological salt response | Ratio of height from the soil to the ligule of the newest expanded leaf in salt over control. Measured on 28 day old plants after 14 days of 9dS salt stress. | Unpublished. Study described in Campbell et al. (2017) |
| Ht FE Ratio (Hfr) | Morphological salt response | Ratio of height (cm) from the soil to the ligule of the newest expanded leaf in salt over control. Measured on 28 day old plants after 14 days of 9dS salt stress. | Unpublished. Study described in Campbell et al. (2017) |
| Panicle number per plant (Pnu) | Yield | Average number of panicles (inflorescences) per plant | Zhao et al. (2011) |
| Panicle length (Pal) | Yield | Length of panicle (inflorescence) from the base to the tip (cm) | Zhao et al. (2011) |
| Primary panicle branch number (Ppn) | Yield | Number of primary branches along the panicle (inflorescence) | Zhao et al. (2011) |
| Seed number per panicle (Snpp) | Yield | Number of seeds per panicle (inflorescence), determined by counting the number of filled spikelets along the main panicle | Zhao et al. (2011) |
| Panicle fertility (Paf) | Yield | Percent of spikelets that filled and produced seeds determined as the ratio of seeds per panicle/spikelets per panicle | Zhao et al. (2011) |
